# Taxonomy, ecology and distribution of *Juniperus oxycedrus* L. group in the Mediterranean Region using morphometric, phytochemical and bioclimatic approaches

**DOI:** 10.1101/459651

**Authors:** Ana Cano Ortiz, Carmelo M. Musarella, José C. Piñar Fuentes, Carlos J. Pinto Gomes, Giovanni Spampinato, Eusebio Cano

## Abstract

The ecology, taxonomy and distribution of the *Juniperus oxycedrus* L. group of taxa are studied. From an ecological aspect, this work proposes a new ombroedaphoxeric index to explain the presence of populations of *Juniperus* in ombrotypes that are not the optimum for these taxa. The controversy among various authors on the taxonomy of the *oxycedrus* group with regard to *J. oxycedrus* subsp. *badia* and *J. oxycedrus* subsp. *lagunae* is clarified. The phytochemical differences in essential oils are addressed, and their similarities analysed; greater similarities are observed between *oxycedrus* and *badia* (H. Gay) Debeaux, and between *navicularis* Grand. and *macrocarpa* (Sm.) Ball. The phytochemical, molecular and distribution differences allow *J. macrocarpa* and *J. navicularis* to be maintained as species.

## Introduction

The prickly juniper species (*Juniperus oxycedrus* L.) is widely distributed throughout the Mediterranean area, and has three subspecies on the Iberian Peninsula: *oxycedrus, badia* and *macrocarpa* [1]. The subspecies *macrocarpa* and *oxycedrus* extend as far as the Balearic Islands, Corsica, Sardinia and the Italian Peninsula [2], whereas the subsp. *oxycedrus* is found as far as Croatia and Slovenia [3]. At one time the *J. oxycedrus* group included *Juniperus navicularis* (syn.: *Juniperus oxycedrus* L. subsp. *transtagana* Franco) and *Juniperus deltoides* R.P. Adams [syn.: *Juniperus oxycedrus* L. subsp. *deltoides* (R.P.Adams) N. G. Passal.], which was described as a new species by [4] for Greece. Subsequently, this same author [5] reported the distribution of *J. deltoides* in Italy, Croatia, Greece and Turkey –coexisting in Turkey with *Juniperus polycarpos* K. Koch–, and established phytochemical differences with *J. oxycedrus* due to its lower alpha-pinene content and higher limonene content. These data have also been confirmed by [6].

All the taxa in the *J. oxycedrus* group grow in xeric environments, generally in inaccessible places such as limestone or siliceous screes, although isolated individuals may appear in *Quercus rotundifolia* Lam. woodlands. The phytocoenoses of *Juniperus* are of considerable ecological interest due to the presence of companion endemics in these plant communities, which serves as the justification for their study. They form small vegetation islands that act as species reservoirs as they are not used for either agriculture or livestock farming and thus have not been destroyed by human action. In these phytocoenoses it is frequent to find endemic species with different degrees of distribution on the peninsula, such as *Echinospartum ibericum* Rivas Mart., Sánchez Mata & Sancho, *Adenocarpus argyrophyllus* (Rivas Goday) Caball., *Digitalis purpurea* L. subsp. *mariana* (Boiss.) Rivas Goday, *Sideritis lacaitae* Font Quer, *Coincya longirostra* (Boiss) Greuter & Burdet, *Cytisus scoparius* (L.) Link subsp. *bourgaei* (Boiss.) Riv.-Mart., *Cytisus striatus* (Hill) Rothm. subsp. *eriocarpus* (Boiss. & Reut.) Rivas-Mart., *Genista polyanthos* R. Roem. ex Willk., *Dianthus crassipes* R. de Roemer, *Dianthus lusitanus* Brot., *Digitalis thapsi* L., *Digitalis purpurea* L. subsp. *heywoodii* P. Silva & M. Silva, *D. purpurea* L. subsp. *mariana* (Boiss) Rivas Goday, *Securinega tinctoria* (L.) Rothm., *Lavandula stoechas* L. subsp. *luisieri* (Rozeira) Rozeira, *Lavandula stoechas* L. subsp. *sampaiana* Rozeira, *Genista hirsuta* Vahl, *Thymus mastichina* L., *Thymus grantensis* Boiss. subsp. *micranthus* (Willk.) O. Bolòs & Vigo, *Thymus zygis* Loefl. ex L. subsp. *gracilis* (Boiss.) R. Morales, *Antirrhinum graniticum* Roth. subsp. *onubensis* (Fernández Casas) Valdés. The territories studied are Sites of Community Interest (SCI) due to their presence on the vertical walls of habitats such as 8210 “Calcareous rocky slopes with chasmophytic vegetation”, and 8220 “Siliceous rocky slopes with chasmophytic vegetation”, which include many endemic plant associations [7] and explain the need to conserve these areas. However, in less steeply sloping rocky areas, the dominant species is *Juniperus oxycedrus* subsp. *badia*, which characterises habitat 5210 “Arborescent matorral with *Juniperus spp.*”. These zones can therefore be classified as hotspots of special interest for conservation. All these associations are included in the Habitats Directive, which justifies the ecological importance of these areas and the need to study them for their subsequent conservation [8].

The areas dominated by *Juniperus* species are currently undergoing a process of expansion due to the increase in areas of scree, which are becoming more widespread each year due to deforestation and forest fire. This phenomenon causes an increase in edaphoxerophilous zones and a decrease in climatophilous zones, thus increasing the potential areas that can act as a refuge for endemic species [9].

The aim of this work is to clarify the taxonomy, ecology and distribution of the taxa in the *J. oxycedrus* group, which form the communities included in the Habitats Directive such as the “Arborescent matorral with *Juniperus spp.*” (5210).

## Materials and Methods

We studied the *J. oxycedrus* group and conducted herborisation and phytosociological sampling campaigns in the field to determine the habitat in which these taxa occur. We examined the differences in the ecology, distribution and taxonomy of the taxa in the *J. oxycedrus* group by analysing their morphological, ecological and phytochemical differences. With these characters, we studied the resemblance between the taxa using a similarity analysis based on the data provided by [10], [11], and [12]. A bioclimatic study was carried out to explain the presence of *Juniperus* species in different rocky substrates.

To understand the presence of communities of *Juniperus* in territories dominated by species in the genus *Quercus*, we applied Thornthwaite’s, ETP_monthly_ = 16(10.T/I)^a^, to calculate the potential evapotranspiration and 0.2ETP by [13]. With these data we used the new ombroedaphoxeric index Ioex proposed by [14] which, in a comparative analysis with the ombrothermic index I_o_ proposed by [15], justifies the presence of microwoodlands of *Juniperus* and *Pinus*.

Territories behave differently in response to the general climate, the type of substrate and the topography of the terrain. For this reason areas on rocky crests –although they may be located in rainy environments and surrounded by climactic forests– behave differently from the territories around them. In these circumstances, islands evolve which may contain edaphoseries, minoriseries and permaseries [16]. All plant communities growing on rocky crests and steeply sloping areas with extreme gradients, among others, are very significantly influenced by the soil, which conditions their existence. All territories have a particular type of substrate and an orography that determines whether they have a greater or lesser capacity to retain water. In ideal situations with good soil texture and structure and with no slopes, the water retention capacity (RC) can be assumed to be maximum (100%). Otherwise, losses occur due to runoff and drainage, causing the RC to vary. Water is also lost through evapotranspiration (ETP). However, as plants have the capacity to self-regulate their losses, it can be accepted that the residual evapotranspiration e = 0.2ETP. There are therefore two parameters (**e** and **RC**) implicated in the development of a vegetation, which is essentially conditioned by rainfall. The ombroclimatic index I_o_ therefore does not explain the presence of plant communities influenced by the substrate, and for this reason, we use the Ombroedaphoxeric Index [14] to explain the presence of communities of *Juniperus* and *Pinus* for territories with a thermotype ranging from the thermo to the supramediterranean.

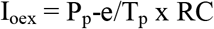

P_p_ = mean annual precipitation. T_p_ = mean annual temperature [15]. e = residual evapotranspiration whose value is 0.2 ETP [13]. RC = retention capacity in parts per unit, whose values may be 0.25, 0.50, 0.75 and 1.

For the authorship of the taxa we have followed [2], [17], [18], [19], [20], [21], [22], [23], [24].

## Results

### Ecology and distribution

*Juniperus oxycedrus* L. has its distribution in the Mediterranean region, eastern Portugal, Morocco and even extends as far as northern Iran [1]. According to this author, this species has three clearly differentiated subspecies. The subspecies *macrocarpa* (Sm.) Ball. is typical of dunes and coastal sand flats, and may occasionally occupy rocky areas. The communities characterized by this taxon on the Iberian Peninsula are described and included in the alliance *Juniperion turbinatae* Rivas-Martínez 1975 *corr*. 1987, along with other plant communities presided by *Juniperus navicularis* Gand. [synonym of *Juniperus oxycedrus* L. subsp. *transtagana* Franco in Feddes Repert. Spec. Nov. Regni Veg. 68:166 (1963)] and *Juniperus phoenicea* L. subsp. *turbinata* (Guss.) Nyman, also typical of psamophilous environments and dunes in coastal zones [25]. The subspecies *macrocarpa* is distributed in Iberian and Italian territories and in the Balearic Islands, Corsica, Sardinia, Sicily (western Mediterranean); this taxon does not reach the eastern Mediterranean and is substituted in Greece by *Juniperus drupacea* Labill..

*J. oxycedrus* L. subsp. *oxycedrus* and *J. oxycedrus* L. subsp. *badia* (H. Gay) Debeaux are found on the Iberian Peninsula on hard acid and basic substrates. The area of distribution of *J. oxycedrus* extends as far as Italy [2], Croatia and Slovenia [3]. It is generally found within *Quercus rotundifolia* Lam. and *Quercus ilex* L. woodlands, and also frequently forms plant communities on screes; as both the subsp. *oxycedrus* and the subsp. *badia* have their optimum development on skeletic-rocky substrates and in semiarid-dry bioclimatic environments (Fig 1).

**Fig 1.**
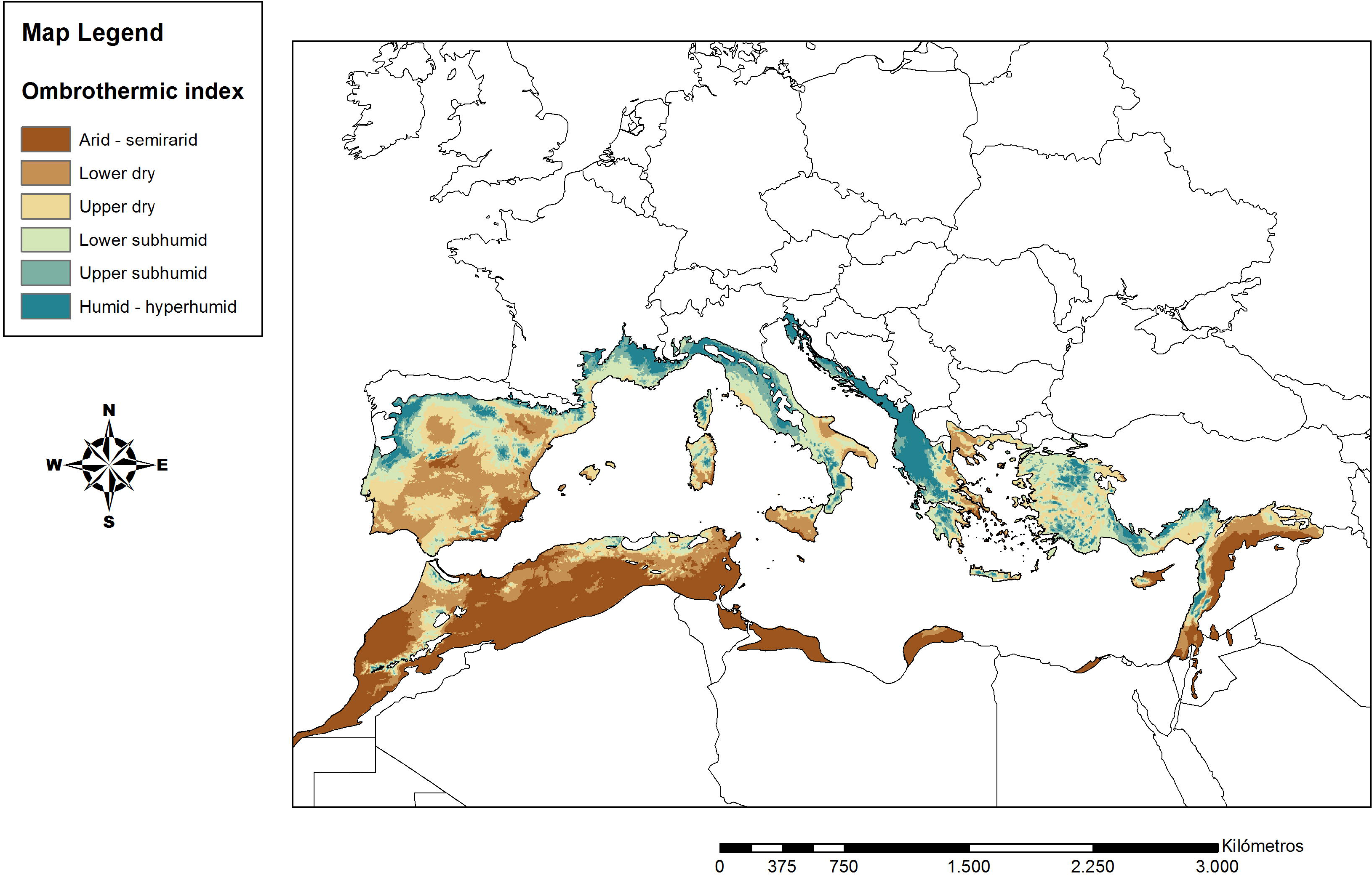
Optimum areas for the development of *Juniperus oxycedrus* group. Optimum ombrotypes: semiarid, dry.

There are also some doubts as to the presence of *J. oxycedrus* subsp. *badia* on the African continent. Some authors such as [26] do not recognise this taxon in northern Africa, although [27] reports *J. oxycedrus* f. *badia* H. Gay in North Africa, and the subsp. *oxycedrus* and subsp. *macrocarpa*.

Bolòs & Vigo [28] include the var. *laguna* Pau *ex* Bolòs et Vigo in *J. oxycedrus* subsp. *oxycedrus*, as it has the same characters as the subspecies *badia*. Recently, based on the work of [20], [29] formulated the new combination *Juniperus oxycedrus* L. subsp. *lagunae* (Pau *ex* C. Vicioso) Rivas Mart. This all serves to highlight the complexity of this taxon, whose distribution area is insufficiently known. However, its presence is very evident in the centre and south of the Iberian Peninsula, where it grows in formations with a broad extension, generally on screes and biotopes with shallow soils where holm oaks (*Quercus rotundifolia*) cease to be dominant or simply cannot exist due to absence of the necessary ecological and/or soil conditions for these taxa to develop [30] (Fig 2).

**Fig 2.**
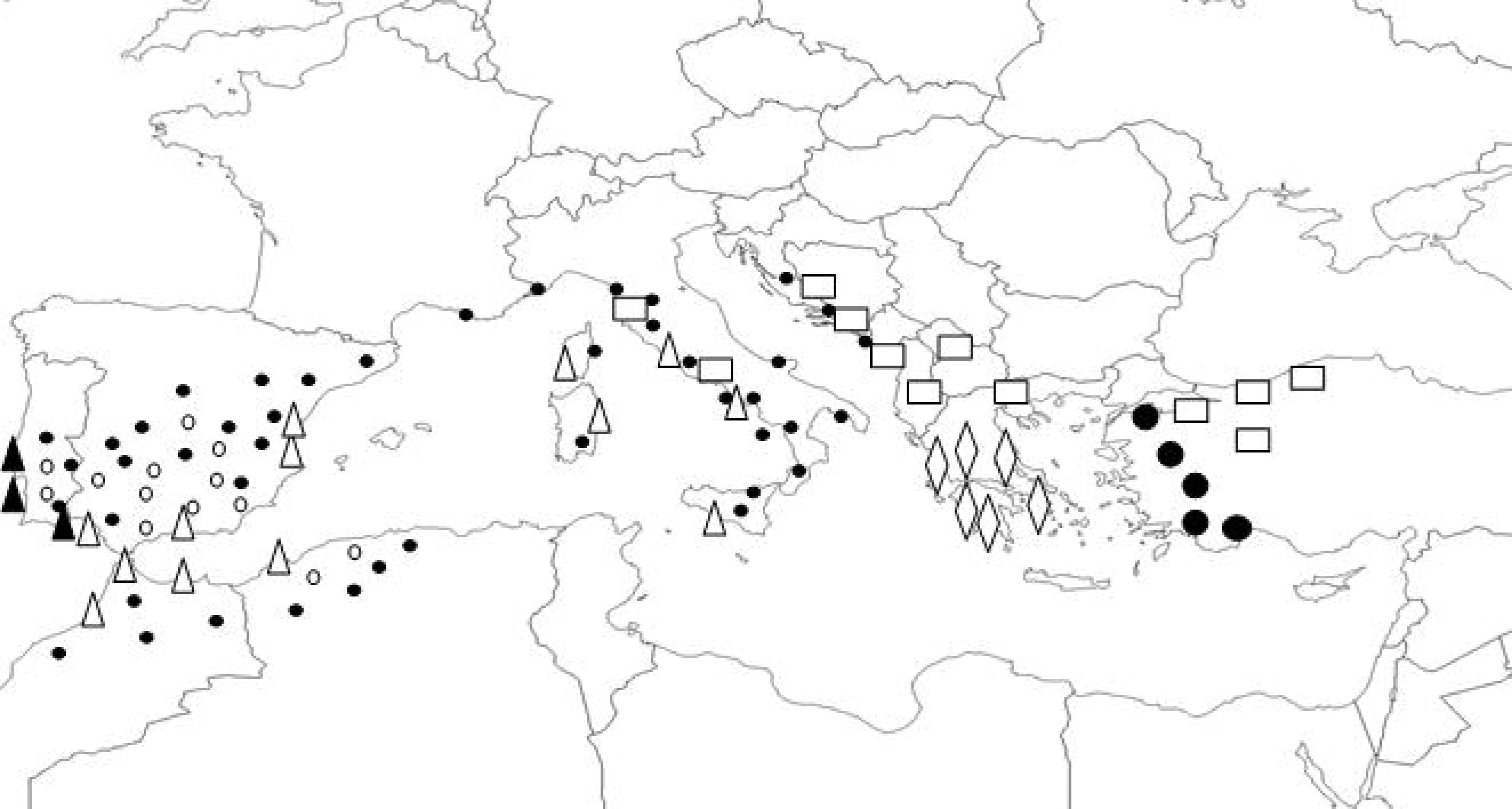
Distribution of taxa of *Juniperus oxycedrus* group in the Mediterranean bassin. • *J. oxycedrus* subsp. *oxycedrus*, ⚬ *J. oxycedrus* subsp. *badia*, ▲ *J. navicularis*, △ *J. macrocarpa*, ⚫ *J. polycarpos*, ▭ *J. deltoides*, ◊ *J. drupacea*.

Table 1 shows the values of Pp, Tp and Io according to the criterion established by Rivas-Martínez. The formula of Thornthwaite ETP_monthly_ = 16(10.T/I)^a^ is applied to obtain the value of ETP, where T is the mean monthly temperature, I is the annual heat index and a parameter that depends on the values taken by I.

**Table 1.** Comparative value of indices Io and I_oex_ in some localities in the southern Iberian Peninsula.

| Weather Station | Pp | Tp | Io | Ombrotype | ETP | e | Ioex1 | Ioex2 | Ioex3 | Ombroclimatic behavior of the locality |
| --- | --- | --- | --- | --- | --- | --- | --- | --- | --- | --- |
| Almadén-Minas (CR) | 625.20 | 194.40 | 3.20 | dry | 808.54 | 161.70 | 0.59 | 1.19 | 1.78 | semiarid |
| Cabezas Rubias (H) | 993.40 | 177.60 | 5.60 | subhumid | 702.14 | 140.42 | 1.20 | 2.40 | 3.60 | dry |
| Aracena (H) | 1025.80 | 175.20 | 5.90 | subhumid | 703.46 | 140.69 | 1.26 | 2.52 | 3.78 | dry |
| Santiago Pontones (J) | 1148.70 | 164.40 | 7.00 | subhumid | 675.23 | 135.04 | 1.54 | 3.08 | 4.62 | dry |
| Vadillo Castril (J) | 1182.20 | 140.40 | 8.40 | humid | 488.88 | 97.72 | 1.93 | 3.86 | 5.79 | subhumid |
| Grazalema (Ca) | 1962.20 | 183.60 | 11.00 | humid | 726.22 | 145.24 | 2.47 | 4.94 | 7.42 | subhumid |
| Montoro (Co) | 522.40 | 210.00 | 2.50 | dry | 903.15 | 180.63 | 0.40 | 0.81 | 1.22 | arid |
| Pozoblanco (Co) | 514.40 | 193.20 | 2.70 | dry | 805.45 | 161.09 | 0.45 | 0.91 | 1.37 | arid |
| Villanueva del Arzobispo (J) | 698.20 | 196.80 | 3.50 | dry | 915.70 | 183.14 | 0.65 | 1.30 | 1.96 | semiarid |

Applying the formula I_oex_ for the assumptions that RC is 0.25, 0.50, 0.75, we obtain three values, of which the most representative is I_oex2._ Table 1 shows the equivalence values so that although the territorial bioclimate allows the existence of climactic forests, in wild areas with RC = 50% the humid ombrotype becomes dry or subhumid depending whether the value of RC =25% or 50%. The subhumid becomes dry and the dry becomes semiarid or arid. Therefore, in areas with Io > 8 we obtain I_oex2_ values of 3.86 and 4.94, which is equivalent to subhumid. This explains the fact that in rocky areas there is an edaphoxerophilous community of *Quercus faginea* s.l. or *Abies pinsapo* Boiss., as occurs in Grazalema (Cadiz); or that a value of I_oex1_ = 2.47 is obtained in the case that RC = 25%. In this situation, there is a presence of an edaphoxerophilous community of *Quercus rotundifolia* in Cazorla (Jaén) and in Grazalema (Cadiz). When the underlying ombrotype is subhumid, the equivalence value of I_oex2_ corresponds to dry; and a starting situation with a dry Io gives semiarid and even arid values of I_oex2_ if we start from a horizon that is lower than dry. This does not allow development of tree species of *Quercus*, but does allow development of the genera *Juniperus* and *Pinus.* The value of I_oex_ is affected by climate change, as evidenced by Del Río et al. [31], who report a heterogeneous trend in terms of annual rainfall redistribution, with a decline in most of the mountainous areas of Grazalema, Ronda, Cazorla, Segura, Sierra Nevada and a large part of the Sierra Morena. However, these authors have detected an increase in rainfall on the Andalusian coast and particularly in Almería. This affects the forest stands, and in conjunction with human activity [8], favours a redistribution of the current forests, with a decline in *Quercus* woodlands and an increase in the micro-woodlands of *Juniperus*.

### Taxonomy

As specified by [10] and [12] there is a clear phytochemical differentiation between the three subspecies of *Juniperus oxycedrus*. Subsequently [5] clarifies the phytochemical differences between *J. oxycedrus* and *J. deltoides*, a species described by this author [4] in Greek territories. This taxon has a low content in alpha-pinene and a high content in limonene, which, among other morphological differences, justifies the rank of species for this taxon. There are also substantial differences in the essential oils of *Juniperus navicularis* Gand. compared to the rest [11].

Alpha-pinene is common to the four taxa and p-cymene to *J. oxycedrus* subsp. *oxycedrus*, *J. oxycedrus* subsp. *macrocarpa* and *J. oxycedrus* subsp. *navicularis* (Table 1). This is justification for including them all in the *Juniperus oxycedrus* group. Moreover, there are certain morphological and ecological similarities between *J. oxycedrus* subsp. *oxycedrus* and *J. oxycedrus* subsp. *badia* and between *J. oxycedrus* subsp. *macrocarpa* and *J. navicularis*, according to the presence of certain oils. The component myrcene is exclusive to subspecies *oxycedrus*, whereas the exclusive compounds in subspecies *badia* are: germacrene D, alpha-camphelenal, beta bourbenene and nine sesquiterpenes. The subspspecies *macrocarpa* does not have any of its own essential oils as it shares ganma-terpinene and terpinen 4-ol with *J. navicularis*, while the exclusive chemical compounds in *J. navicularis* are alpha-phellandrene, alpha-terpinene, terpinolene, alpha terpinol and ten sesquiterpenes.

The Pearson correlation analysis (Tables 2 and 3, Fig 3) gives values of r = 1 or near 1 for group G2 (*J. oxycedrus* subsp. *macrocarpa* and *J. navicularis*). In the case of group G1 (*J. oxycedrus* subsp. *oxycedrus* and *J. oxycedrus* subsp. *badia*) the values of rare close or equal to 1; however the relation between both groups is low owing to the phytochemical differences between them.

**Table 2.**
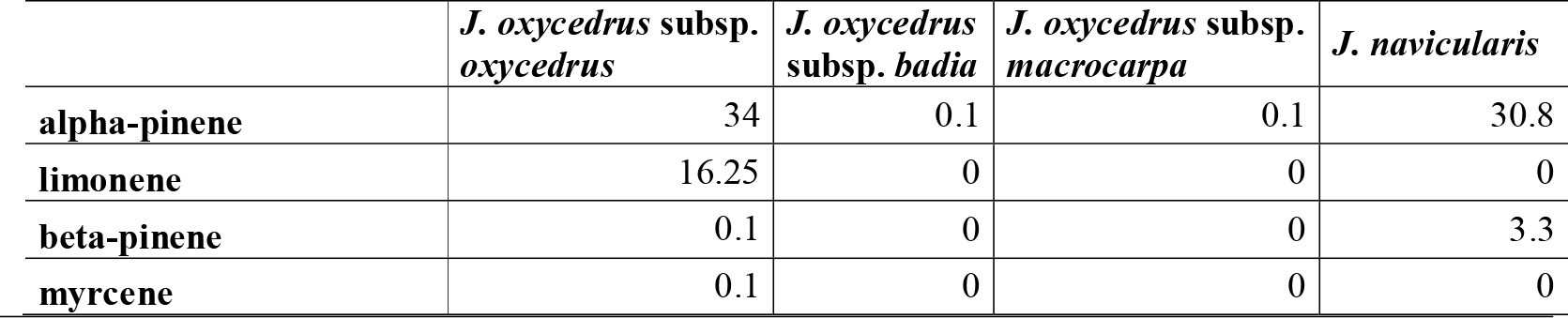
Essential oil content in four species of *Juniperus* from [10, 11].

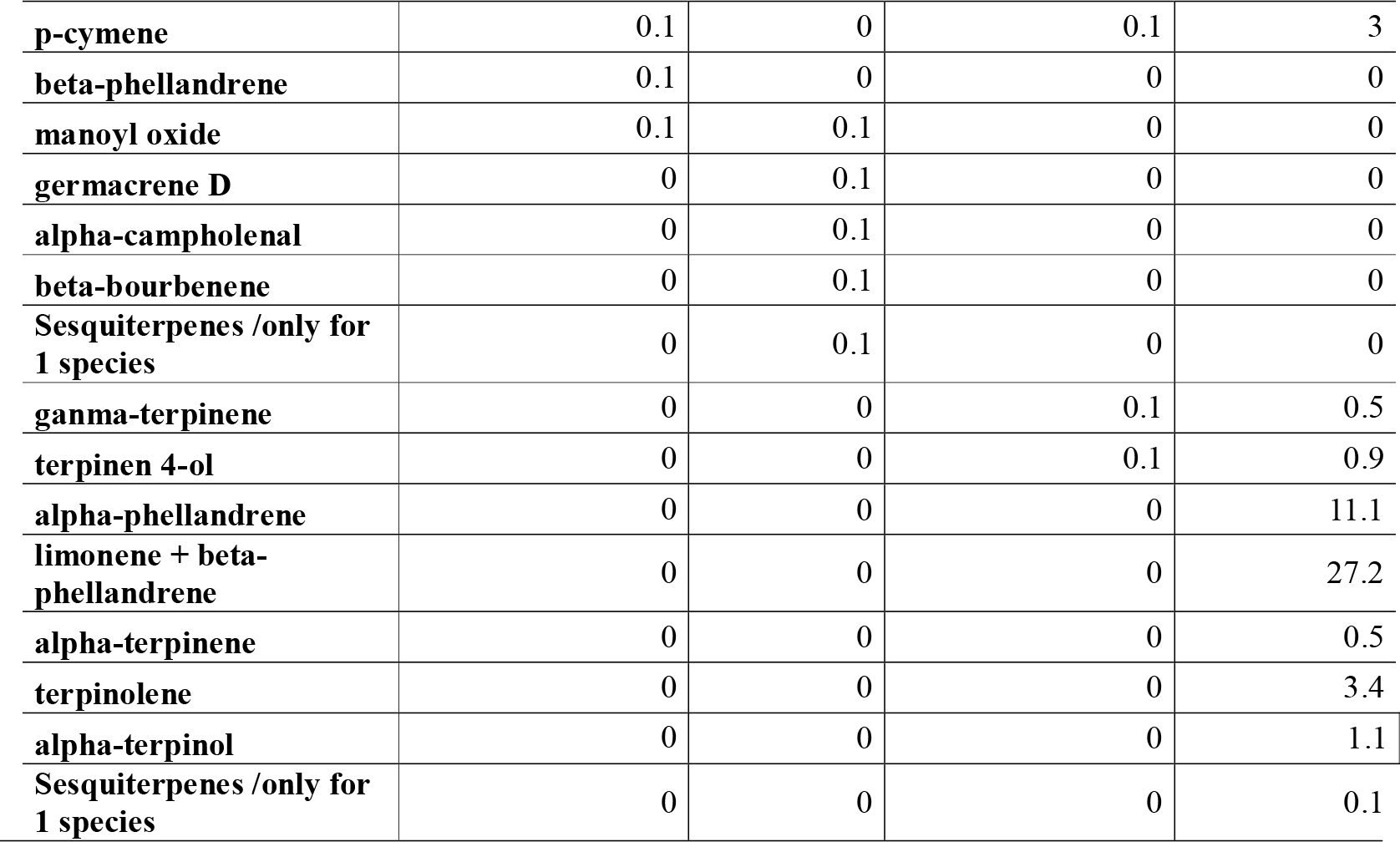

**Table 3.** r- Pearson. The values in bold are different from 0 with a level of significance of alpha=0.05.

|  | 1 | 2 | 3 | 4 | 5 | 6 | 7 | 8 | 9 | 10 | 11 | 12 | 13 | 14 | 15 | 16 | 17 | 18 | 19 |
| --- | --- | --- | --- | --- | --- | --- | --- | --- | --- | --- | --- | --- | --- | --- | --- | --- | --- | --- | --- |
| 1 | 1 | 0,633 | 0,543 | 0,633 | 0,532 | 0,633 | 0,049 | -0,576 | -0,576 | -0,576 | -0,576 | 0,424 | 0,470 | 0,519 | 0,519 | 0,519 | 0,519 | 0,519 | 0,519 |
| 2 | 0,633 | 1 | -0,306 | **1,000** | -0,318 | **1,000** | 0,577 | -0,333 | -0,333 | -0,333 | -0,333 | -0,420 | -0,382 | -0,333 | -0,333 | -0,333 | -0,333 | -0,333 | -0,333 |
| 3 | 0,543 | -0,306 | 1 | -0,306 | **1,000** | -0,306 | -0,565 | -0,347 | -0,347 | -0,347 | -0,347 | **0,977** | **0,992** | **1,000** | **1,000** | **1,000** | **1,000** | **1,000** | **1,000** |
| 4 | 0,633 | **1,000** | -0,306 | 1 | -0,318 | **1,000** | 0,577 | -0,333 | -0,333 | -0,333 | -0,333 | -0,420 | -0,382 | -0,333 | -0,333 | -0,333 | -0,333 | -0,333 | -0,333 |
| 5 | 0,532 | -0,318 | **1,000** | -0,318 | 1 | -0,318 | -0,590 | -0,363 | -0,363 | -0,363 | -0,363 | **0,983** | **0,995** | **0,999** | **0,999** | **0,999** | **0,999** | **0,999** | **0,999** |
| 6 | 0,633 | **1,000** | -0,306 | **1,000** | -0,318 | 1 | 0,577 | -0,333 | -0,333 | -0,333 | -0,333 | -0,420 | -0,382 | -0,333 | -0,333 | -0,333 | -0,333 | -0,333 | -0,333 |
| 7 | 0,049 | 0,577 | -0,565 | 0,577 | -0,590 | 0,577 | 1 | 0,577 | 0,577 | 0,577 | 0,577 | -0,728 | -0,662 | -0,577 | -0,577 | -0,577 | -0,577 | -0,577 | -0,577 |
| 8 | -0,576 | -0,333 | -0,347 | -0,333 | -0,363 | -0,333 | 0,577 | 1 | **1,000** | **1,000** | **1,000** | -0,420 | -0,382 | -0,333 | -0,333 | -0,333 | -0,333 | -0,333 | -0,333 |
| 9 | -0,576 | -0,333 | -0,347 | -0,333 | -0,363 | -0,333 | 0,577 | **1,000** | 1 | **1,000** | **1,000** | -0,420 | -0,382 | -0,333 | -0,333 | -0,333 | -0,333 | -0,333 | -0,333 |
| 10 | -0,576 | -0,333 | -0,347 | -0,333 | -0,363 | -0,333 | 0,577 | **1,000** | **1,000** | 1 | **1,000** | -0,420 | -0,382 | -0,333 | -0,333 | -0,333 | -0,333 | -0,333 | -0,333 |
| 11 | -0,576 | -0,333 | -0,347 | -0,333 | -0,363 | -0,333 | 0,577 | **1,000** | **1,000** | **1,000** | 1 | -0,420 | -0,382 | -0,333 | -0,333 | -0,333 | -0,333 | -0,333 | -0,333 |
| 12 | 0,424 | -0,420 | **0,977** | -0,420 | **0,983** | -0,420 | -0,728 | -0,420 | -0,420 | -0,420 | -0,420 | 1 | **0,996** | **0,980** | **0,980** | **0,980** | **0,980** | **0,980** | **0,980** |
| 13 | 0,470 | -0,382 | **0,992** | -0,382 | **0,995** | -0,382 | -0,662 | -0,382 | -0,382 | -0,382 | -0,382 | **0,996** | 1 | **0,994** | **0,994** | **0,994** | **0,994** | **0,994** | **0,994** |
| 14 | 0,519 | -0,333 | **1,000** | -0,333 | **0,999** | -0,333 | -0,577 | -0,333 | -0,333 | -0,333 | -0,333 | **0,980** | **0,994** | 1 | **1,000** | **1,000** | **1,000** | **1,000** | **1,000** |
| 15 | 0,519 | -0,333 | **1,000** | -0,333 | **0,999** | -0,333 | -0,577 | -0,333 | -0,333 | -0,333 | -0,333 | **0,980** | **0,994** | **1,000** | 1 | **1,000** | **1,000** | **1,000** | **1,000** |
| 16 | 0,519 | -0,333 | **1,000** | -0,333 | **0,999** | -0,333 | -0,577 | -0,333 | -0,333 | -0,333 | -0,333 | **0,980** | **0,994** | **1,000** | **1,000** | 1 | **1,000** | **1,000** | **1,000** |
| 17 | 0,519 | -0,333 | **1,000** | -0,333 | **0,999** | -0,333 | -0,577 | -0,333 | -0,333 | -0,333 | -0,333 | **0,980** | **0,994** | **1,000** | **1,000** | **1,000** | 1 | **1,000** | **1,000** |
| 18 | 0,519 | -0,333 | **1,000** | -0,333 | **0,999** | -0,333 | -0,577 | -0,333 | -0,333 | -0,333 | -0,333 | **0,980** | **0,994** | **1,000** | **1,000** | **1,000** | **1,000** | 1 | **1,000** |
| 19 | 0,519 | -0,333 | **1,000** | -0,333 | **0,999** | -0,333 | -0,577 | -0,333 | -0,333 | -0,333 | -0,333 | **0,980** | **0,994** | **1,000** | **1,000** | **1,000** | **1,000** | **1,000** | 1 |

Types of oils studied: 1. Alpha-pinene, 2. Limonene, 3. Beta-pinene, 4. Myrcene, 5. P-cymene, 6. Beta-phellandrene, 7. Monoyl oxide, 8. Germacrene D, 9. Alpha-campholenal, 10. Beta-bourbenene, 11. Sesquiterpenes exclusives, 12. Gamma-terpinene, 13. Terpinen 4-ol, 14 Alpha-phellandrene, 15. Limonene + beta-phellandrene, 16. Alpha-terpinene, 17. Terpinolene, 18. Alpha-terpinol, 19. Sesquiterpenes.

**Fig 3.**
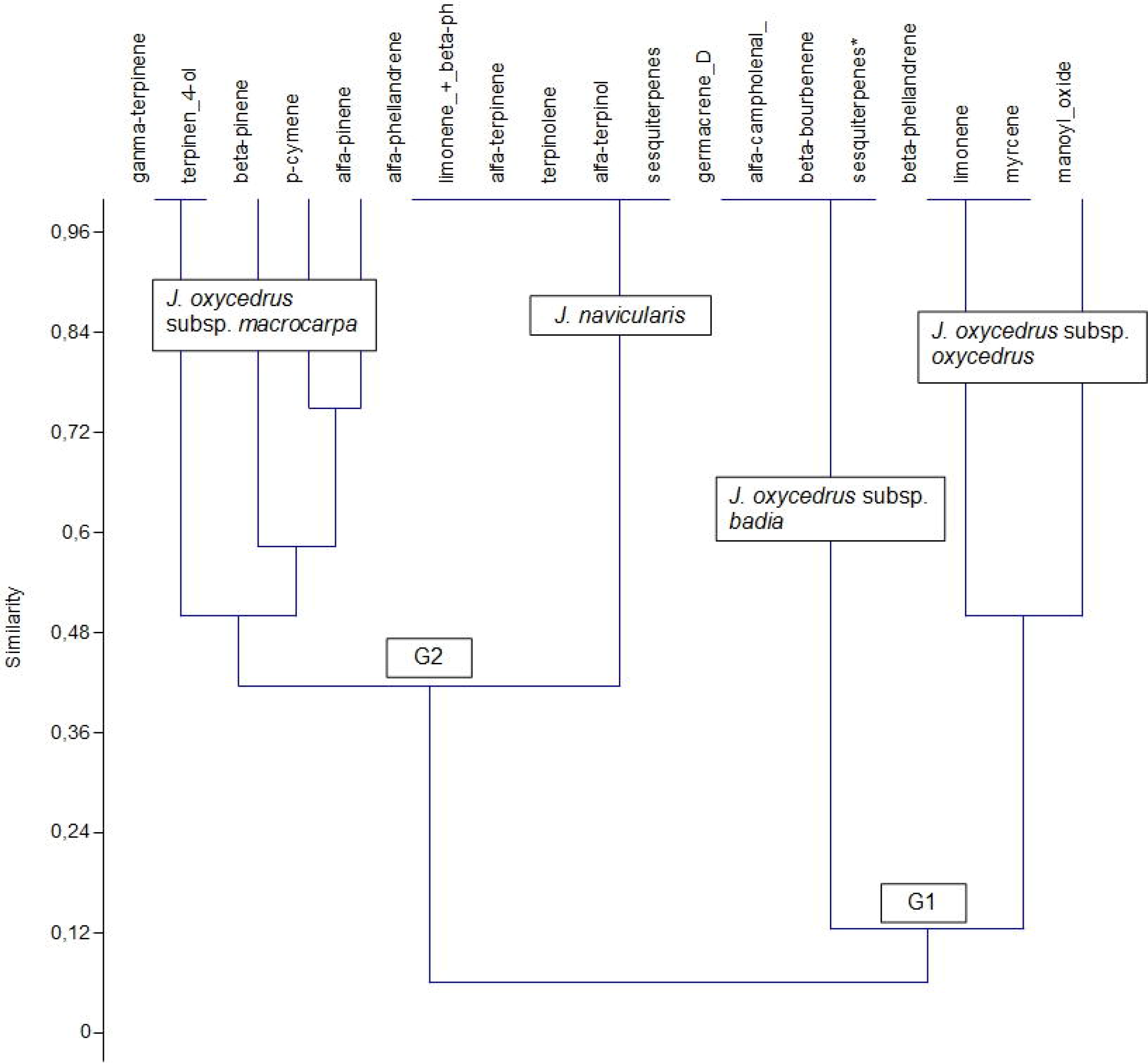
Similarity analysis between the taxa *Juniperus oxycedrus* subsp.*oxycedrus*, subsp.*badia*,*J. oxycedrus* subsp.*macrocarpa* and *J*.*navicularis.*

## Discussion

Linnaeus in the first part of his work *Speics Plantarum* did not describe *Juniperus oxycedrus* L., where he merely noted the existing synonymy. It was Clusius who later described this species and gave specific localities for Spain, which were subsequently included by [32].

The main differences between *Juniperus oxycedrus* subsp. *oxycedrus* and *Juniperus oxycedrus* subsp. *badia* are based on their physiognomy and the size of their mature fruits [1]. Whereas the subspecies *oxycedrus* tends to occur as a shrub, the subspecies *badia* (H. Gay) Debeaux is a tree of considerable height and with a pyramidal form. The size of the mature galbuli in the subspecies *oxycedrus* does not generally exceed 1 cm, while in the subspecies *badia* it is over 1 cm. The leaves of *oxycedrus* have a width of 1-1.5 mm while those of subsp. *badia* are 1.2-2 mm. Coincidentally these subspecies are frequently found coexisting in similar biotopes, which has led to frequent confusion by several authors. The subsp. *macrocarpa* (Sm.) Ball. has galbuli of 1.2-1.5 cm. with a purplish chestnut colour when ripe; this is an erect tree, rarely prostrate, growing up to 3 m. In contrast *Juniperus navicularis* Gand. has a similar ecology to subsp. *macrocarpa* but grows to a maximum height of 2 m; its leaves are 1-1.5 mm wide, and its galbuli are between 0.7-1 cm (Table 4).

**Table 4.** Morphometric comparison in the taxa of *Juniperus oxycedrus* group from [1].

|  | *J. oxycedrus*<br>subsp. *oxycedrus* | *J. oxycedrus*<br>subsp. *badia* | *J. oxycedrus* subsp.<br>*macrocarpa* | *J.*<br>*navicularis* |
| --- | --- | --- | --- | --- |
| **leaf length (mm)** | 8-15(25) mm | 12-20 mm | 20-25 mm | 4-12 mm |
| **leaf width (mm)** | 1-1.5 mm | 1.2-2 mm | 2-2.5 mm | 1-1.5 mm |
| **fruit (gabuli)** | 8-10 mm | 10-13 mm | 12-15 mm | 7-10 mm |

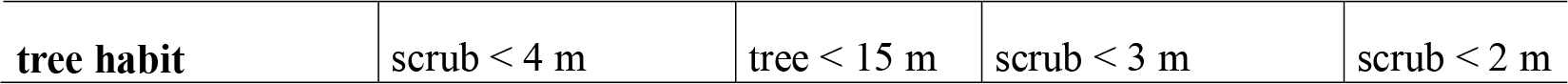

Theoretically, the authorship of *Juniperus oxycedrus* L. should be attributed to Clusius, who described it: *“Nusquam autem majorem vidisse memini, quam supra Segoviam et Guadarrama, itinere Madritiano, ubi magnarum arborum interdum altitudinem”.* He also says of the fruit *“fructum initio viridem... postremo, cum maturuit punicea colorís, Juniperi fructu multo majorem ut interdum avellanam magnitudine aequet”* [33]. This description is specifically included in the work of [19]. Clusius’ descrtiption, in which he states literally that it has a “thick pruinose red fruit”, corresponds to *Juniperus oxycedrus* L. subsp. *badia* (H. Gay) Debeaux [Basión. *Juniperus oxycedrus* var. *badia* H. Gay in Assoc. Franç. Avancem. Sci. Compt. Rend. 1889:501 (1889)], included in [1]. The lack of typification of the specimen described by Clusius is justification for these facts, and supports the valid acceptance of *Juniperus oxycedrus* L. subsp. *badia* (H. Gay) Debeaux (ICN article 41). Due to the high number of morphological and ecological differences and in essential oils between subsp. *macrocarpa*, subsp. *oxycedrus* and subsp. *badia*, and to its similarity with *J. navicularis*, the subspecies *macrocarpa* can be proposed to be subordinated to *J. navicularis* as *J. navicularis* Gand. subsp. *macrocarpa* (Sm.) Ball; however, the molecular study by [34] establishes major molecular differences between the four taxa *J. oxycedrus* subsp. *oxycedrus, J. oxycedrus* subsp. *badia, J. oxycedrus* subsp. *macrocarpa* and *J. navicularis*, and based on this he raises them to the rank of species. The phytochemical differences between groups G1 and G2 are very clear, so the species rank could be accepted for the four taxa. However, in the case of their distribution, the populations of *J. oxycedrus* subsp. *macrocarpa* and *J. navicularis* are separated despite having similar ecologies, while this is not the case of *J. oxycedrus* subsp. *oxycedrus* and subsp. *badia*. Therefore, it is logical to accept *J. macrocarpa* Sm. and *J. navicularis* Gand. as species and to maintain *J. oxycedrus* L. subsp. *oxycedrus* and *J. oxycedrus* L. subsp. *badia* (H. Gay) Debeaux with the rank of subspecies.

Finally, considering the phytosociological point of view, in vegetation studies the subspecies *badia* (H. Gay) Debeaux and the subspecies *lagunae* (Pau ex Vicioso) Rivas-Martínez have been used indistinctly as a result of accumulated and persistent errors [30]; the name *Juniperus oxycedrus* L. subsp. *lagunae* (Pau ex Vicioso) Rivas-Martínez is invalid and based on the iconography of Laguna by Pau [*Juniperus oxycedrus* L. subsp. *lagunae* (Pau ex C.Vicioso) Rivas Mart. - Itinera Geobot. 15(2): 703 (2002); nom. inval.], as the name of the basionym is not validly published (ICN articulo 7). The correct name is therefore *Juniperus oxycedrus* L. subsp. *badia* (H. Gay) Debeaux. Based on these facts, the various authors of plant associations must rectify their names.

## Conclusions

In this work, we update the state of knowledge on the ecology, taxonomy and distribution of the taxa of the *Juniperus oxycedrus* group, based on our study and on research by various authors, which has occasionally been the source of controversy. The similarity study based on the quantity, type of essential oils, molecular analyses and different bioclimatic distribution justify maintaining the rank of species for *J. macrocarpa* Sm. and *J. navicularis* Gand.. Instead, the distribution and co-existence of the two remaining taxa, in spite of their phytochemical and molecular differences, do not warrant the rank of species, and we thus maintain their rank as subspecies: *J. oxycedrus* L. subsp. *oxycedrus* and *J. oxycedrus* L. subsp. *badia* (H. Gay) Debeaux. This study, in accordance to others [6, 35, 36, e.g.] also confirm that for a good discrimination among species of groups with difficult interpretation, morphometric, phytochemical and bioclimatic approaches are very useful to clarify the belonging rank to the taxa. All these results and conclusions are according to [37].

## Acknowledgments

Ms Pru Brooke-Turner (M.A. Cantab.), a native English speaker specialized in scientific texts, has translated this article. This manuscript has been released as a Pre-Print at: Biorxiv 459651; https://doi.org/10.1101/459651 2018.

## Author Contributions

**Conceptualization**: Eusebio Cano.

**Data curation**: Eusebio Cano; Ana Cano Ortiz; Carmelo M. Musarella; José C. Piñar Fuentes; Carlos J. Pinto Gomes; Giovanni Spampinato.

**Formal analysis**: Ana Cano Ortiz; Carmelo M. Musarella; José C. Piñar Fuentes.

**Investigation**: Eusebio Cano; Ana Cano Ortiz; Carmelo M. Musarella; José C. Piñar Fuentes; Carlos J. Pinto Gomes; Giovanni Spampinato.

**Methodology**: Eusebio Cano; Ana Cano Ortiz.

**Resources**: Eusebio Cano; Ana Cano Ortiz; Carmelo M. Musarella; Carlos J. Pinto Gomes; Giovanni Spampinato.

**Supervision**: Eusebio Cano; Carmelo M. Musarella.

**Validation**: Eusebio Cano.

**Visualization**: Eusebio Cano; Ana Cano Ortiz.

**Writing ± original draft**: Eusebio Cano; Ana Cano Ortiz; Carmelo M. Musarella; José C. Piñar Fuentes; Carlos J. Pinto Gomes; Giovanni Spampinato.

**Writing ± review & editing**: Eusebio Cano; Ana Cano Ortiz; Carmelo M. Musarella.

